# Separable functions of the PHD finger protein Spp1 in the Set1 and the meiotic DSB forming complexes cooperate for meiotic DSB formation

**DOI:** 10.1101/181180

**Authors:** Céline Adam, Raphaël Guérois, Anna Citarella, Laura Verardi, Florine Adolphe, Claire Béneut, Vérane Sommermeyer, Claire Ramus, Jérôme Govin, Yohann Couté, Valérie Borde

## Abstract

Histone H3K4 methylation is a feature of meiotic recombination hotspots shared by many organisms including plants and mammals. Meiotic recombination is initiated by programmed double-strand break (DSB) formation that in budding yeast takes place in gene promoters and is promoted by histone H3K4 di/trimethylation. This histone modification is recognized by Spp1, a PHD-finger containing protein that belongs to the conserved histone H3K4 methyltransferase Set1 complex. During meiosis, Spp1 binds H3K4me3 and interacts with a DSB protein, Mer2, to promote DSB formation close to gene promoters. How Set1 complex- and Mer2- related functions of Spp1 are connected is not clear. Here, combining genome-wide localization analyses, biochemical approaches and the use of separation of function mutants, we show that Spp1 is present within two distinct complexes in meiotic cells, the Set1 and the Mer2 complexes. Disrupting the Spp1-Set1 interaction mildly decreases H3K4me3 levels and does not affect meiotic recombination initiation. Conversely, the Spp1-Mer2 interaction is required for normal meiotic recombination initiation, but dispensable for Set1 complex-mediated histone H3K4 methylation. Finally, we evidence that Spp1 preserves normal H3K4me3 levels independently of the Set1 complex. We propose a model where the three populations of Spp1 work sequentially to promote recombination initiation: first by depositing histone H3K4 methylation (Set1 complex), next by “reading” and protecting histone H3K4 methylation, and finally by making the link with the chromosome axis (Mer2-Spp1 complex). This work deciphers the precise roles of Spp1 in meiotic recombination and opens perspectives to study its functions in other organisms where H3K4me3 is also present at recombination hotspots.

**Author summary:** Meiotic recombination is a conserved pathway of sexual reproduction that is required to faithfully segregate homologous chromosomes and produce viable gametes. Recombination events between homologous chromosomes are triggered by the programmed formation of DNA breaks, which occur preferentially at places called hotspots. In many organisms, these hotspots are located close to a particular chromatin modification, the methylation of lysine 4 of histone H3 (H3K4me3). It was previously shown in the budding yeast model that one protein, Spp1, plays an important function in this process. We further explored the functional link between Spp1 and its interacting partners, and show that Spp1 shows genetically separable functions, by depositing the H3K4me3 mark on the chromatin, “reading” and protecting it, and linking it to the recombination proteins. We provide evidence that Spp1 is in three independent complexes to perform these functions. This work opens perspectives for understanding the process in other eukaryotes such as mammals, where most of the proteins involved are conserved.

## Introduction

In all reproducing organisms, recombination between homologous chromosomes at meiosis is essential to form viable gametes with normal chromosome content. Meiotic recombination is initiated by the programmed formation of DSBs catalyzed by the conserved Spo11 protein together with largely conserved accessory DSB proteins (1-3) and is required to promote genetic diversity and for accurate homolog segregation (4). Histone modifications are key players of the chromatin organization. Among these, histone H3K4 trimethylation (H3K4me3) is able to recruit downstream effectors such as chromatin remodelers. In meiosis, Set1, the subunit of the Set1 complex that catalyzes histone H3K4 methylation, is required for normal levels and distribution of meiotic double-strand breaks (5, 6). In budding yeast, the vast majority of meiotic DSBs occur in the c.a. 140 bp nucleosome depleted regions (NDR) at gene promoters, close to nucleosomes bearing the histone H3K4me3 modification (5, 7). The tendency of meiotic recombination to occur in gene promoters and H3K4me3 regions is conserved among many organisms such as dogs, plants and birds (8-10). In mice and humans, meiotic recombination also occurs at H3K4me3 sites, but only those deposited by the meiotic-specific histone H3K4 methyltransferase, PRDM9 (11-13). PRDM9 recognizes specific DNA sequences with its array of zinc finger motifs, where it methylates H3K4 and promotes recombination, away from gene promoters (14). Remarkably, the mutation of *Prdm9* in mice redirects meiotic recombination events towards gene promoters and H3K4me3 (15), as if PRDM9 had a dominant role over the default promoter/histone H3K4me3 conserved pathway.

The link between histone H3K4 methylation and meiotic DSB formation has recently been explained in budding yeast by the role of the PHD finger protein, Spp1, in spatially linking DSB sites to the recombination initiation machinery (16, 17). During meiotic prophase, chromosomes adopt a specific three-dimensional structure formed of chromatin loops anchored at their basis to a chromosome axis (18). DSB are formed into loop DNA sequences, whereas the DSB proteins are located on the chromosome axis, implying a spatial contact between these two physically distant chromosomal regions during DSB formation (3, 19-21). In meiosis, Spp1, a member of the Set1 complex, is, like Set1, required for normal DSB levels. Spp1 is specifically important for H3K4 trimethylation, and in its absence, H3K4me3 levels are reduced to about 20% of wild-type (17). This has been attributed to Spp1 being important for opening the catalytic site of Set1 and allowing trimethylation (22). Spp1 also interacts with Mer2, one of the axis-associated DSB proteins required for DSB formation, and is preferentially located on the chromosome axis. The PHD finger of Spp1 interacts also with H3K4me2/me3 at +1 nucleosomes and is required for normal DSB formation, and thus Spp1 make the physical link between gene promoters close to H3K4me2/3 sites and the DSB formation machinery (16, 17). Thus, Spp1 may facilitate or stabilize the interaction between these distant regions, which triggers DSB formation by Spo11, the protein that bears the catalytic DSB forming activity (16, 17).

In vegetative cells, Spp1 belongs to the Set1 complex and its distribution mirrors that of RNA pol II (17). By contrast, in meiosis, the chromosomal distribution of Spp1 shows no spatial correlation with that of RNA pol II (17), raising the question whether Spp1 is still part of the Set1 complex in meiosis, and if so, how it distributes between the Set1 complex and the DSB proteins. In addition, given the role of Spp1 for H3K4me3, known to recruit downstream chromatin remodelers, it is not clear as well if the functional role of Spp1 within the Set1 complex for H3K4 methylation can be separated from its implication in DSB formation through its interaction with Mer2.

In this paper, we evidenced that Spp1 interacts both with the Set1 complex and Mer2 in meiotic cells. However, Set1 does not associate with chromosomes axes in meiosis, and its subunits do not interact with Mer2, revealing that Spp1 is present in two distinct complexes. Next, we show that surprisingly, the presence of Spp1 in the Set1 complex is not important for maintaining H3K4 trimethylation levels and that Spp1 acts independently of the Set1 complex to promote meiotic DSB formation. Finally, we show that a mutant of Mer2 that no longer interacts with Spp1 but binds normally to chromosome axes is impaired for DSB formation. This demonstrates that solely affecting Spp1 interaction with Mer2 is sufficient to impair DSB formation, independently of any H3K4 methylation-related change in chromatin. Finally this work is relevant for understanding meiotic DSB formation in mammals and other organisms, for which a mechanism linking H3K4me3 and the DSB machinery likely exists but has not yet been elucidated.

## Results

### Spp1 is associated with both the DSB protein Mer2 and the Set1 complex in meiosis

During vegetative growth, Spp1 is a member of the eight-unit Set1 complex, and predominantly locates at highly transcribed genes, consistent with the Set1 complex being associated with elongating RNA pol II (17, 23). However, in meiosis, Spp1 interacts with the Mer2 protein and is predominantly located on chromosomes axes (17). We thus investigated how Spp1 distributes between the Set1 complex and its interaction with the axis-associated Mer2 DSB protein in meiotic cells. For this, we affinity purified Spp1-TAP from cells at 3.5 hr in meiotic prophase, the expected time of DSB formation. The Spp1-TAP fusion protein associated with chromosome axis sites during meiosis and is thus fully functional (Fig 1A). Spp1-TAP interacting proteins were purified and identified by label-free mass-spectrometry-based quantitative proteomics. Analyses revealed that Spp1-TAP interacts with the whole Set1 complex and with Mer2 in meiotic cells (Fig 1B and S1 Table). No other protein was significantly purified with Spp1.

**Fig 1.**
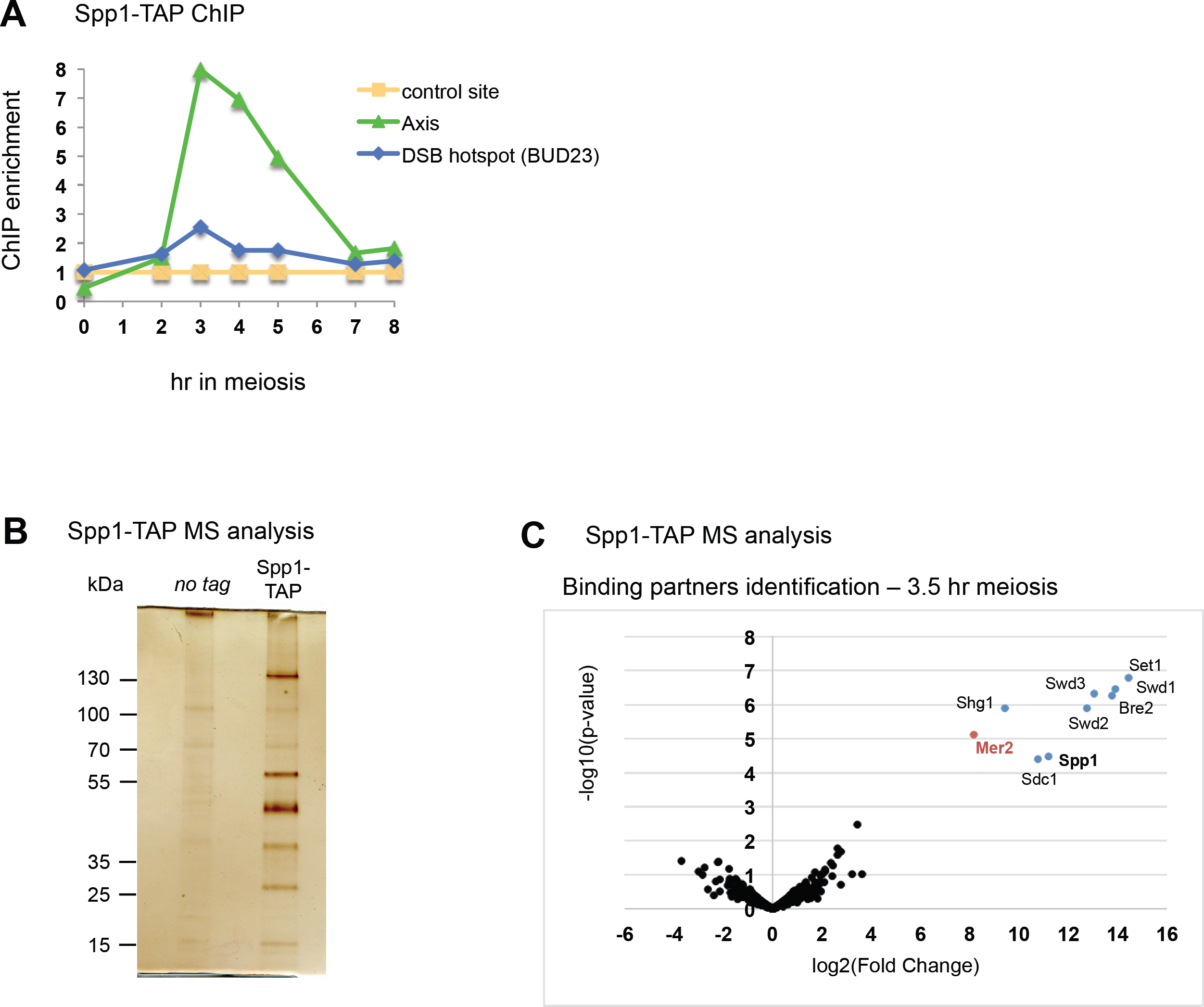
Spp1 interacts with the whole Set1 complex and the Mer2 DSB protein in meiotic cells. (A) ChIP-qPCR of Spp1-TAP during meiosis, showing its association with the chromosome axis at 3-4 hr, the expected time for DSB formation. Strain: VBD1266. (B) Silver-stained gel of TAP-purification eluates performed at 3.5 hr in meiosis in an untagged (ORD7339) or Spp1-TAP (VBD1266). (C) Mass spectrometry analysis of proteins pull-down by Spp1-TAP in meiosis (t = 3.5 hr). Spp1-TAP: VBD1266 – no tag: ORD7339. The volcano plot indicates in color the proteins significantly co-purified with Spp1 (log2(Fold change)>3 and log10(p-value)<4). The experiment was done in triplicate.

### Distinct requirements govern Spp1 association with highly transcribed genes and with chromosome axis in meiotic cells

Our finding that Spp1 is associated with the Set1 complex in meiosis is in apparent contradiction with the fact that Spp1 appears mainly associated with regions of axis attachment, largely co-localizing with Mer2, as assessed by genome wide mapping (17). The contribution of Mer2 to the localization of Spp1 was assessed by mapping genome-wide Spp1 binding sites in *mer2Δ* meiotic cells. Whereas in wild-type, Spp1 binding was not correlated with that of RNA pol II, in *mer2Δ*, Spp1 location became positively correlated with that of RNA pol II (Pcorr=0.57) (Fig 2A). Indeed, in the absence of Mer2, the strongest Spp1 peaks were now at highly transcribed genes, similar to the distribution of Spp1 in vegetative cells, likely reflecting its association with the Set1 complex (Fig 2B). The localization of Spp1 was confirmed by ChIP-qPCR in WT and *mer2Δ* cells. Interestingly, in wild-type cells, we were able to detect Spp1 at two individual highly transcribed genes, *ACS1* and *CIT2*, like in *mer2Δ*, in addition to its association with chromosome axes (Fig 2D). By contrast, Spp1-Myc fully associated with meiotic chromosomes axes in the absence of Set1 (Fig 2C and S1 Fig), but the signal at the two highly transcribed genes *ACS1* and *CIT2* was lost (Fig 2D).

**Fig 2.**
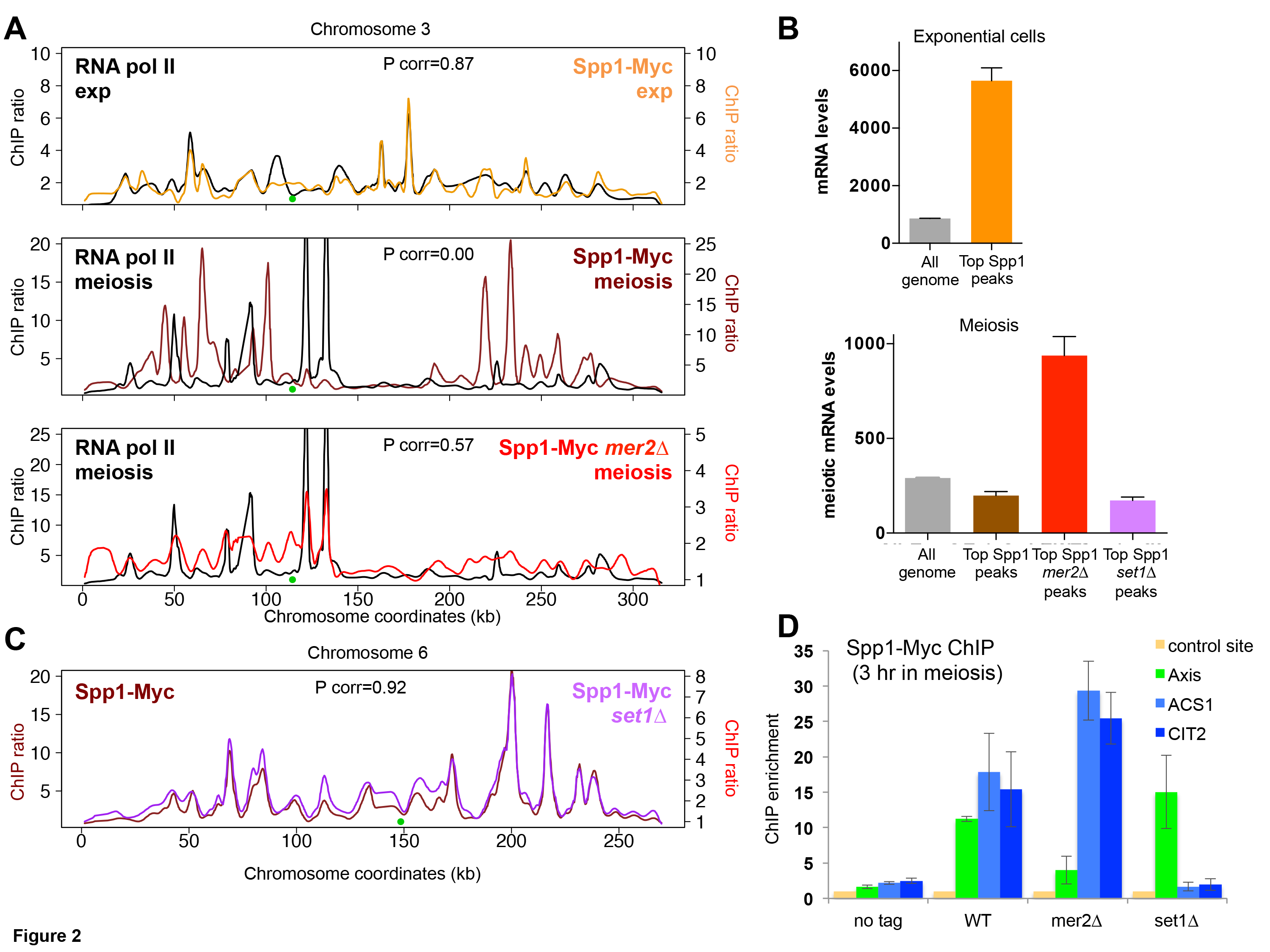
Spp1 binds distinct Set1- and Mer2-dependent sites in meiotic cells. (A) Comparison of Spp1 binding profile with that of RNA polymerase II, in exponential cells, or in meiosis (t = 3 hr), in wild-type or *mer2Δ* mutant. Spp1-Myc and RNA pol II binding data are from (17). Spp1-Myc *mer2Δ*: VBD1220. ChIPchip profiles are shown for chromosome 3. Centromere is indicated as a green dot, and ratios are plotted after smoothing with a 2 kb window. (B) Mean mRNA levels at the 100 strongest Spp1 peaks or in the whole genome. Spp1 peaks were determined from the experiments shown in A. mRNA levels are from published data in SK1 diploid exponential cells (54) (upper panel) or SK1 meiotic cells at t = 4 hr (55) (lower panel). Error bars: S.E.M. (C) Spp1 binding profile in *SET1* or *set1Δ* cells in meiosis (t = 3 hr). Spp1-Myc *set1Δ*: VBD1209. ChIPchip profiles are shown for chromosome 6. Centromere is indicated as a green dot, and ratios are plotted after smoothing with a 2 kb window. (D) qPCR analysis of Spp1 binding to an axis site and two highly transcribed genes in meiosis, *ACS1* and *CIT2* (t = 3 hr). No tag: ORD7339; WT: VBD1187; *mer2Δ*: VBD1220; *set1Δ*: VBD1209. Values are mean ± S.E.M. of two independent experiments.

We propose that in wild-type cells, the association of Spp1 with Mer2 occurs at the highly localized axis association sites, resulting in strong peaks that mask its association with chromatin through the Set1 complex, which may be more diffuse along chromatin, and less concentrated in strongly defined peaks. Our findings are consistent with the existence of two classes of Spp1 binding sites, ones with axis sites, dependent on Mer2, and ones with transcription sites, dependent on Set1.

### Set1 does not bind chromosome axes during meiosis and the Set1 complex does not associate with Mer2

We next asked if when Spp1 is associated with Mer2 on chromosomes axes, it is within the Set1 complex. To answer this, we mapped the sites of chromatin association of Set1 in meiosis. Set1 bound the two highly transcribed genes, *ACS1* and *CIT2*, during meiosis, but only weakly a chromosome axis-associated site (Fig 3A and S2 Fig). This was confirmed genome-wide, where the location of Set1 in meiotic cells was positively correlated with that of RNA pol II and Set1 bound only weakly to chromosome axis sites (Figs 3B and 3C, respectively). Consistent with these findings, the strongest Set1 peaks in the genome were at highly transcribed regions (Fig 3D). Genome-wide mapping and individual locus analysis also revealed that Set1 associates with centromeres, contrary to Spp1 (Fig 3C and S2 Fig). This may be a consequence of Set1 acting at kinetochores to methylate the non-histone Dam1 protein (24).

**Fig 3.**
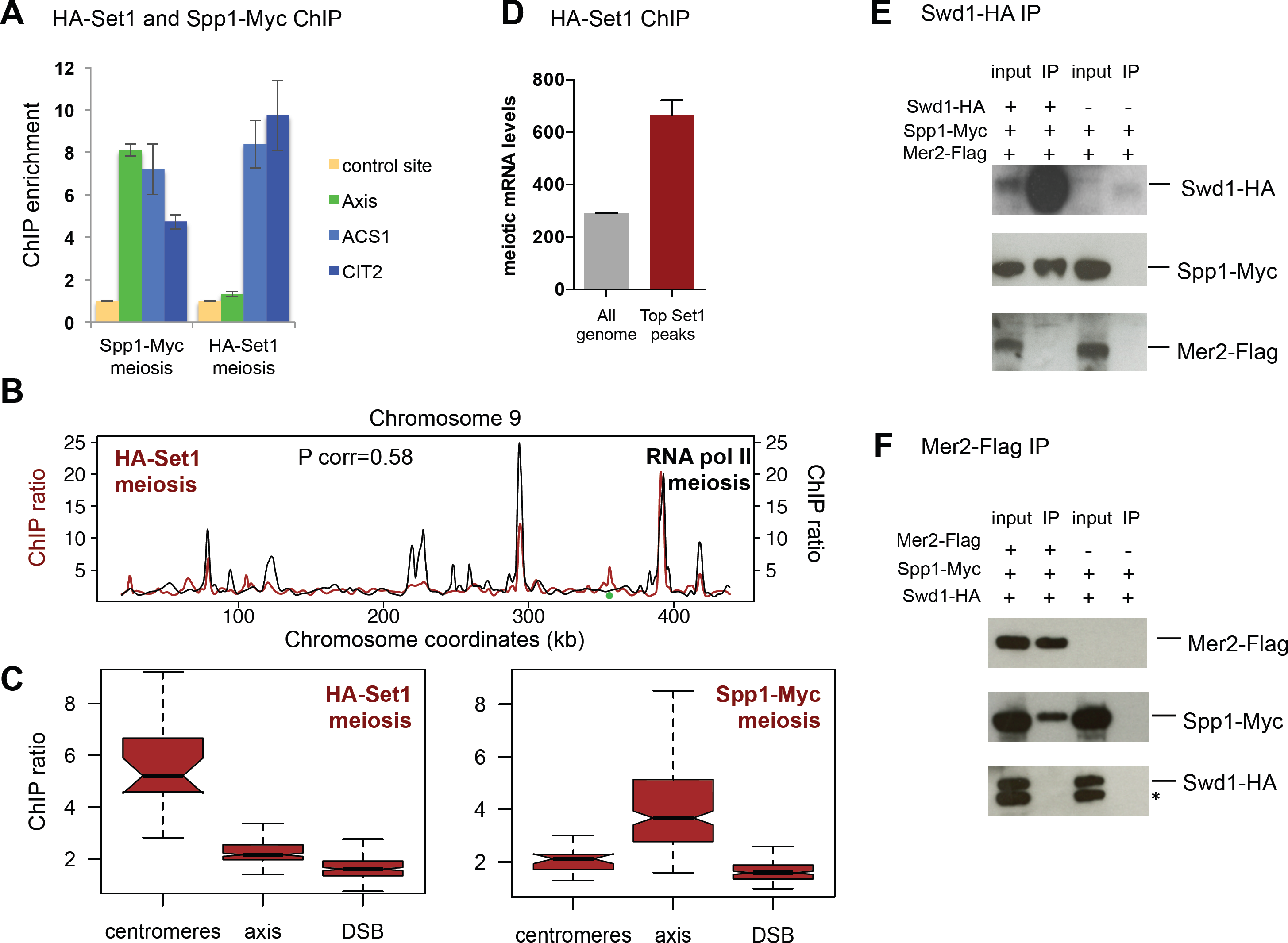
Set1 associates in meiosis with centromeres and highly transcribed genes, but not with chromosome axis sites. (A) ChIP-qPCR of Set1 (t =4 hr) and Spp1 (t = 3 hr) during meiosis comparing their association with axis and two highly transcribed genes. HA-Set1: VBD1378; Spp1-Myc: VBD1187. See also S2 Fig. (B) Chromosome profiles of Set1 and RNA Pol II binding in meiosis. HA-Set1: strain VBD1378 at t = 4 hr. RNA Pol II: data are from (17). (C) ChIPchip signal at the indicated features. For centromeres, mean signal at probes less than 200bp from a centromere; axis and DSB: mean signal at the 200 strongest Red1 and DSB peaks, respectively, as defined in (53). Boxplots indicate median (line), 25th–75th percentile (box) 61.5 times the interquartile range (whiskers). Non-overlapping notches of two boxes are indicative that medians are statistically different. (D) mean mRNA levels at the 100 strongest Set1 peaks or in the whole genome. mRNA levels are from SK1 diploid meiotic cells at t = 4 hr (55). Error bars: S.E.M. (E) Co-immunoprecipitation by the core Set1 complex Swd1 protein from cells at 3.5 hr in meiosis analyzed by western blot. Swd1-HA Spp1-Myc Mer2-Flag: VBD1401; Spp1-Myc Mer2-Flag: VBD1395. (F) Co-immunoprecipitation by the DSB protein Mer2 from cells at 3.5 hr in meiosis. Swd1-HA Spp1-Myc Mer2-Flag: VBD1401; Spp1-Myc Swd1-HA: VBD1400. The asterisk indicates non-specific cross-hybridizing band.

In agreement with our genome-wide localization results, Swd1, a subunit of the Set1 complex, immunoprecipitated Spp1 from meiotic cells, but not Mer2, and reciprocally, Mer2 pulled down Spp1, but not Swd1 (Figs 3E and 3F, respectively). The latter result was confirmed by mass spectrometry-based analysis of Mer2-TAP interactome (Table S1). Together, our experiments point toward Spp1 being located in two physically distinct complexes.

### A mutant that disrupts the Set1-Spp1 interaction reveals that Spp1 works independently of the Set1 complex to promote DSB formation

We next set out to determine if the presence of Spp1 in the Set1 complex is required for its function within the DSB formation complex. For this, we designed a mutation to disrupt the Set1-Spp1 interaction without affecting Spp1’s PHD finger or the interaction of Spp1 with Mer2.

The domain of Spp1 that interacts with Set1 has not been determined, and we failed to identify a Spp1 mutant that would disrupt its interaction with Set1. On the Set1 side, a short region (aminoacids 762-794) close to the regulatory nSet domain of Set1 is sufficient for the interaction with Spp1 in a two-hybrid test (25). We searched for conserved aminoacids in this region that could potentially be involved in protein-protein interactions (26). We identified a conserved negatively charged acidic motif (AIKDEEDM) that we mutated into a neutral motif (ASKSSSSM) (*set1_sid* mutant for <u>S</u>pp1 interaction-<u>d</u>eficient) (Fig 4A and S3 Fig). Remarkably, the mutated Set1 protein lost all interaction with Spp1 in a two-hybrid assay, while keeping its interaction with its two other known direct binding partners, Shg1 and Swd2 (Fig 4B) (27). Furthermore, the *set1_sid* mutation induced the loss of detectable interaction of Spp1 with the Set1 complex *in vivo*, as assessed by the loss of interaction between Spp1 and the Swd1 subunit, both in vegetative and in meiotic cells (Fig 4C) and by mass spectrometry analysis of Spp1-TAP co-purified proteins from meiotic *set1_sid* cells (S4 Fig and S1 Table). Thus, no other subunit of the Set1 complex seems able to retain Spp1 on its own. In addition, in exponential or in meiotic *set1_sid* cells, Spp1-Myc was no longer detected at the *ADH1*, *ACS1* and *CIT2* highly transcribed genes, whereas it still associated with chromosome axis in meiotic cells (Fig 4D). This *set1_sid* mutant allowed us to assess if DSB formation occurs normally even if Spp1 is not associated with the Set1 complex. Remarkably, meiotic DSB frequency at two Spo11 DSB hotspots, *CYS3* and *DEP1*, in the *set1_sid* mutant was indistinguishable from wild-type (Fig 4E, upper panel). In situations where the interaction between H3K4 methylation and Spp1 is disrupted, such as in the *set1Δ* and *spp1Δ* mutants, although DSB frequency is generally reduced, DSB formation is increased at a few sites, including the *PES4* gene promoter (5, 17) Importantly, in the *set1_sid* mutant, no DSB was induced at the *PES4* promoter (Fig 4E, lower panel). These results clearly indicate that the presence of Spp1 in the Set1 complex is dispensable for DSB formation.

**Fig 4.**
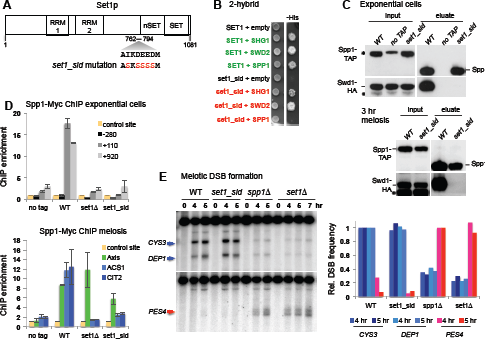
A mutant in an acidic patch of Set1 abolishes its interaction with Spp1 but does not affect meiotic DSB formation. (A) Scheme of the Set1 protein domains, with the Spp1 interacting domain (762-794) (25), RRM1 and RRM2 RNA recognizing motifs (57), the SET catalytic and nSET regulatory domains (58) and the mutations creating the *set1_sid* mutant. (B) 2-hybrid assays for the interaction between Mer2 and Spp1. Growth on the – His medium indicates an interaction between the two tested proteins. (C) Interaction of Spp1 with the Set1 complex subunit Swd1 in exponential or meiotic cells examined by Western blot as indicated. Proteins pulled down by Spp1-TAP were released in the eluate after Tev cleavage of the Tap tag. WT: VBD1745; no TAP: VBD1742; *set1_sid*: VBD1836. The asterisk indicates non-specific cross-hybridizing band. (D) Binding of Spp1 detected by ChIP-qPCR in exponential cells or in meiotic cells at t = 3 hr. No tag: ORD7339; WT: VBD1187; *set1Δ*: VBD1209; *set1_sid*: VBD1868. Values are the mean ± S.E.M. of two independent experiments (E) Meiotic DSB formation monitored in *dmc1Δ* cells by Southern blot at *CYS3* and *DEP1* DSB (upper panel), or at the *spp1Δ*-specific *PES4* DSB (lower panel). DSB sites are indicated by an arrow. WT: ORD7354; *set1_sid*: VBD1854; *spp1Δ*: VBD1748; *set1Δ*: ORD9624. Graph shows the DSB quantification relative to the level in WT (for *CYS3*, *DEP1*) or *spp1Δ* cells (for *PES4* site). DSB were quantified at the 5 hr time point, with the additional 7 hr time point for *set1Δ*.

### Spp1 maintains H3K4me3 levels independently of the Set1 complex

Since Spp1 is believed to facilitate the Set1 catalysis of H3K4 trimethylation, we would expect that in the *set1_sid* mutant where Spp1 is no longer associated with the Set1 complex, H3K4me3 levels should decrease similarly to *spp1Δ* mutant, to about 20% of WT levels ((17) and Figs 5A and 5B). Surprisingly, the *set1_sid* mutant still showed high levels of H3K4me3, as assessed by Western blot and chromatin immunoprecipitation (Figs 5A and 5B). Furthermore, H3K4me3 levels were reduced in the double *set1_sid spp1Δ* compared to the single *set1_sid* mutant, indicating that Spp1 still promotes H3K4me3 levels outside of the Set1 complex (Figs 5A and 5B). One possibility is that its binding to H3K4me3 stabilizes this mark and/or protects it from active demethylation. We asked if this function could be through its PHD finger, known to recognize H3K4 methylation *in vitro* (28). A deletion of the PHD finger or the point mutation *spp1W45A* of the PHD finger only mildly affects global levels of H3K4me3 (16) and Fig 5A). However, a *set1_sid spp1W45A* double mutant showed a reduction of H3K4me3 similar to that of the *set1_sid spp1Δ* mutant (Figs 5A and 5B). This uncovers an unprecedented role of the PHD finger of Spp1 in the maintenance of H3K4me3 levels.

**Fig 5.**
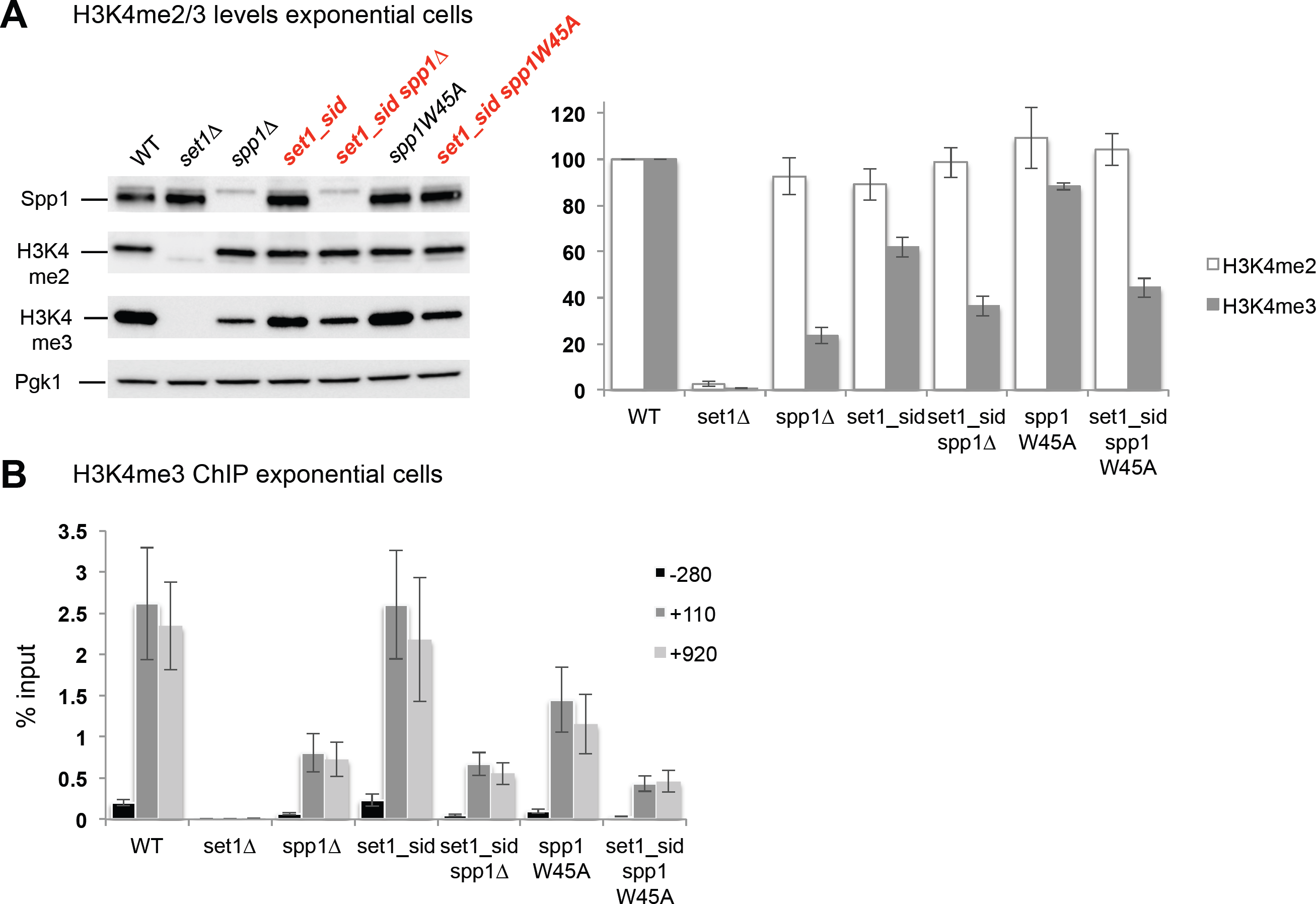
In the *set1_sid* mutant, Spp1 is still important to maintain H3K4me3 levels. (A) Histone H3K4 methylation levels in exponentially growing cells detected by Western blot. WT: ORT4601; *set1Δ*: ORT4784; *spp1Δ*: VBH152; *set1_sid*: VBH1881; *set1_sid spp1Δ*: VBH1972; *spp1W45A*: VBH1419; *set1_sid spp1W45A*: VBH2021. Values underneath each panel indicate protein levels normalized to Pgk1 levels and relative to the WT strain. Values are the mean ± S.E.M. of the normalized relative levels from 2 to 6 replicates for each strain. (B) Histone H3K4me3 levels in exponentially growing cells detected by ChIP at the highly transcribed *ADH1* gene. Same strains as in E. Values are expressed as % of input DNA, and are the mean ± S.E.M. of six independent experiments.

### A Mer2 mutant that no longer interacts with Spp1 mimics *spp1Δ* for DSB formation

Likewise, we asked if the *spp1Δ* phenotype for DSB formation is solely due to absence of Spp1 within the Mer2 complex, and not to indirect effects on Set1 complex function. For this, we designed a mutant that would disrupt the Mer2-Spp1 interaction but preserves the Spp1-Set1 interaction. The domain of Spp1 interacting with Mer2 lies in the last 131 amino acids of Spp1, and the deletion of four amino acids C263 to C266 in this domain was proposed to abolish Mer2-Spp1 interaction *in vivo* and be important for new DSB targeting by a Gal4BD-Spp1 fusion (16). However, this Spp1 mutant retained two-hybrid interaction with Mer2, and when inserted behind its endogenous promoter, was largely proficient in DSB formation at the tested hotspots (S5 Fig). We thus looked for altering the region of Mer2 that interacts with Spp1, with as little as possible alteration of Mer2 other functions. This region has been mapped to aminoacids 165 to 232 (16), which corresponds to one of the two major coiled coils of Mer2 predicted structure (Fig 6A). Combining analyses of structure prediction and conservation of aminoacids in this region, we identified several conserved aminoacids predicted to be surface exposed (Fig 6A and S6 Fig). The mutation of one of these, V195 to D, totally abolished the two-hybrid interaction of Mer2 with Spp1 (*mer2_sid* mutant, Fig 6B). When tested *in vivo*, the Mer2_sid mutant protein was also totally deficient for interaction with Spp1 in meiotic cells (Fig 6C). We examined the meiotic phenotypes of this Spp1 interaction-defective Mer2 mutant. Remarkably, the *mer2_sid* mutant had reduced DSB formation at the two Spo11 hotspots, *CYS3* and *DEP1*, like *spp1Δ* (Fig 6D, upper panel). In addition, DSB formation at the *PES4* site was detected at increased level in the *mer2_sid* mutant, similarly to *spp1Δ* (Fig 6D, lower panel). We conclude that the *mer2_sid* mutation recapitulates all meiotic DSB phenotypes of *spp1Δ*.

**Fig 6.**
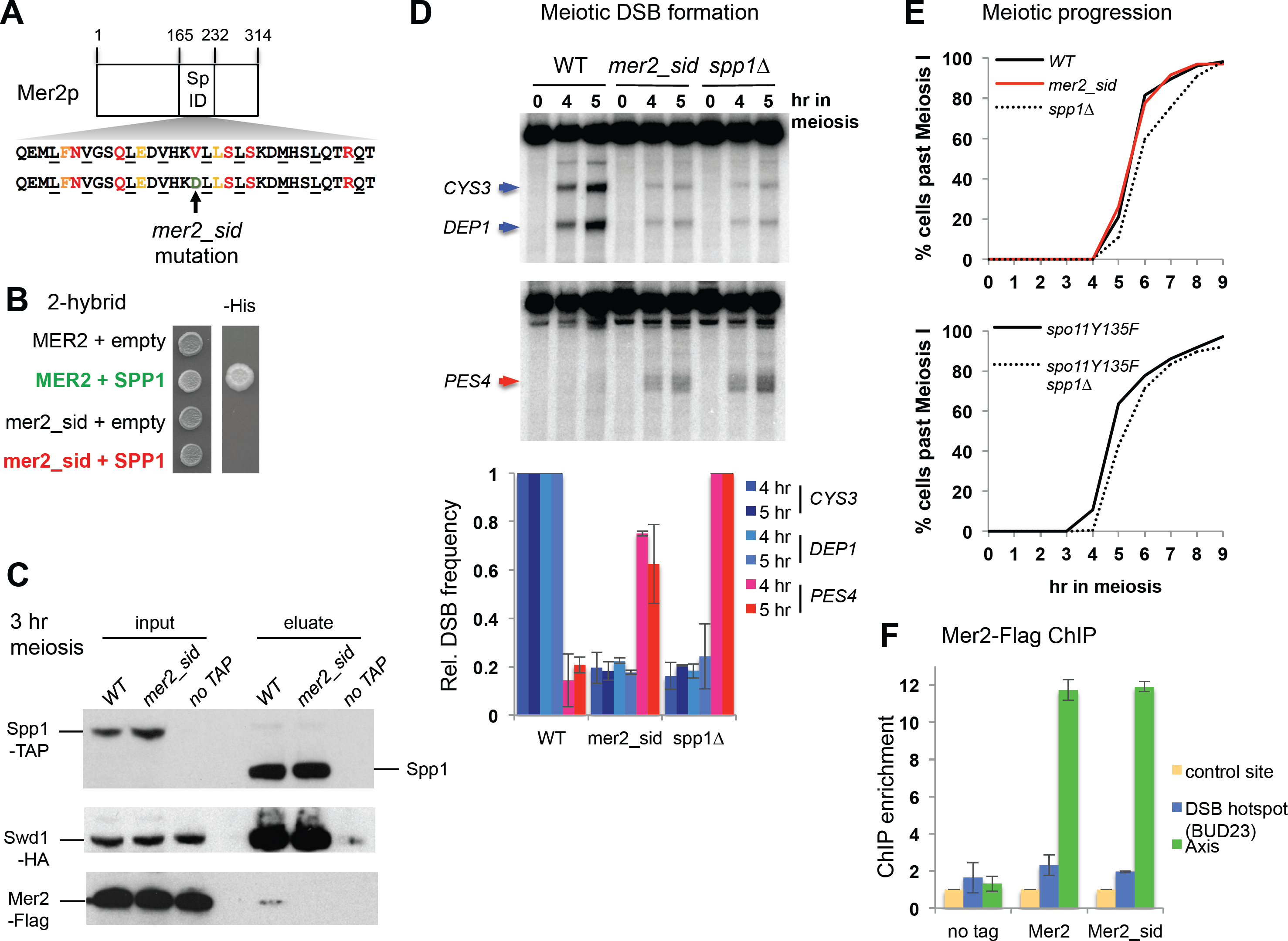
A *mer2* point mutant affected for its interaction with Spp1 mimmics the meiotic phenotype of *spp1Δ*. (A) Scheme of the Mer2 protein, with its predicted coil-coil structures (Hhpred) and the Spp1 interacting domain (165-232) (16). Below is the Mer2 protein sequence in the predicted coiled-coiled heptad structure. Buried positions required for coiled-coil pairing are underlined. Amino acids conserved and predicted to be surface-exposed are colored. The *mer2_sid* mutation (V195D) is indicated by an arrow. (B) 2-hybrid assays for the interaction between Mer2 and Spp1. Growth on the – His medium indicates an interaction between the two tested proteins. (C) Interaction of Spp1 with Mer2 and the Set1 complex subunit Swd1 in meiotic cells examined by Western blot as indicated. Proteins pulled down by Spp1-TAP were released in the eluate after Tev cleavage of the Tap tag. WT: VBD1745; *mer2_sid*: VBD1852; no TAP: VBD1742.(D) Meiotic DSB formation monitored in *dmc1Δ* cells by Southern blot at *CYS3* and *DEP1* DSB (upper panel), or at the *spp1Δ*-specific *PES4* DSB. DSB sites are indicated by an arrow. WT: ORD7354; *mer2_sid*: VBD1879; *spp1Δ*: VBD1748. Graph shows the DSB quantification relative to the level in WT (for *CYS3*, *DEP1*) or *spp1Δ* cells (for *PES4* site). DSB were quantified at the 5 hr time point. Values are the mean ± S.E.M. from two independent experiments. (E) Meiotic progression as assessed by DAPI staining of strains with the indicated genotype. WT: ORD7339; *mer2_sid*: VBD1880; *spp1Δ*: VBD1769; *spo11Y135F*: VBD1291; *spo11Y135F spp1Δ*: VBD1233. (F) Association of Mer2 and Mer2_sid mutant proteins with chromosome axis during meiosis (t = 3 hr) monitored by ChIP. No tag: ORD7339; Mer2-Flag: VBD1251. Mer2_sid Flag: VBD1843.

Next, we checked meiotic progression, since *spp1Δ* was reported to have a meiotic delay (17). The *mer2_sid* mutant had wild-type meiotic progression (Fig 6E, upper panel). The meiotic delay of *spp1Δ* may thus be unrelated to a recombination phenotype of *spp1Δ* but to defects related to the Set1 complex. To answer this question, we examined meiotic progression in a *spo11Y135F* DSB deficient strain. Indeed, *spp1Δ* still showed a meiotic delay in this context, which is thus DSB-independent (Fig 6E, lower panel).

Finally, the *mer2_sid* showed a wild-type level of spore viability (97% among 100 tetrads), contrary to *mer2Δ*, but similar to *spp1Δ* (17). Since Mer2 is an essential DSB protein, this indicates that Mer2-sid keeps its core meiotic functions, apart from its interaction with Spp1. Indeed, Mer2_sid protein was still recruited to the chromosome axis like the non-mutated protein (Fig 6F).

Altogether, these results highlight that all meiotic DSB formation defects of *spp1Δ* can be attributed to its lack of interaction with Mer2, and are unrelated to any change in chromatin opening or accessibility that might be due to affecting the Set1 complex.

## Discussion

We dissected the respective contribution of Spp1 to the Set1 and the DSB formation complexes, and show that they act as independent complexes. We further demonstrate that the meiotic DSB phenotype of *spp1Δ* cells is solely due to the function of Spp1 for tethering DSB sites to the chromosome axis, and not changes in chromatin structure.

### Identification of a Spp1-Mer2 complex that does not comprise other stably associated meiotic DSB proteins

Mer2 is one of the ten DSB proteins identified in *S. cerevisiae*, which are all connected by physical interactions, and was proposed to form a complex with two other DSB proteins, Mei4 and Rec114 (29, 30). However, we did not identify any other candidate than Mer2 and the Set1 complex in the meiotic Spp1 purifications. Likewise, we previously found that Spp1 does not interact with Mei4 in CoIP experiments (17). Furthermore, in a meiotic Mer2-TAP pulldown, we retrieved Spp1 as the top candidate, but no peptide of Mei4 or Rec114 (Table S1). This indicates that the strength or the frequency of interaction between Mer2 and Spp1 is higher than that of Mer2 with Mei4 and Rec114. Spp1 interacts strongly with Mer2 in two-hybrid assays, and thus this interaction does not need *in vivo* modification of Mer2 (16). By contrast, the co-immunoprecipitation of Mei4 and Rec114 with Mer2 and the recruitment of Mei4 and Rec114 to the chromosome axis require Mer2 to be phosphorylated, upon DNA replication, by CDK and DDK (21, 31-33). Thus the interaction of Mer2 with Mei4 and Rec114 is perhaps more transient than the Mer2-Spp1 interaction and too weak to be detected in our analysis. Finally, our *mer2_sid* mutant, despite losing interaction with Spp1, still likely keeps interaction with Mei4 and Rec114, which are essential DSB proteins. We thus propose two separable functions of Mer2: one in recruiting the essential DSB proteins Mei4 and Rec114, and the other in increasing the tethering of DSB sites to the axis, through Spp1, in order to favor cleavage by Spo11. The first function is regulated by replication and Mer2 phosphorylation, whereas the latter is a “constitutive” function of Mer2.

### Spp1 plays a role outside the Set1 complex for maintaining H3K4me3 levels

Histone H3K4me3 levels are slightly decreased in the *set1_sid* mutant, consistent with the hypothesis that the presence of Spp1 in the Set1 complex is important for normal H3K4me3 levels. However, it is surprising that histone H3K4me3 levels were much less affected in the *set1-sid* mutant, than in the *spp1Δ* mutant, revealing a function of Spp1 for histone H3K4me3 levels in the absence of detectable interaction with the Set1 complex. This function thus cannot be attributed to its proposed stimulation of Set1 catalytic activity for H3K4me3 deposition (22). The *in vivo* function of Spp1 recognition of H3K4me3 by its PHD finger has been poorly studied. Within the Set1 complex, it may restrict Set1 activity to the +1 nucleosome of genes, which harbor the combination of H3K4me2/3 and H3R2 not asymmetrically methylated that is specifically recognized by Spp1 PHD finger (34). However, this explanation does not hold for the effect we saw in the *set1_sid* mutant. We thus propose that the PHD finger module of Spp1 binds and protects H3K4me3 from demethylation by the Jhd2/JARID1 enzyme, which specifically demethylates H3K4me3 *in vivo* (35, 36) (Figure 7). Further studies will be required to test this hypothesis and the crosstalk between Spp1 and Jhd2 through their binding to H3K4me3.

**Fig 7.**
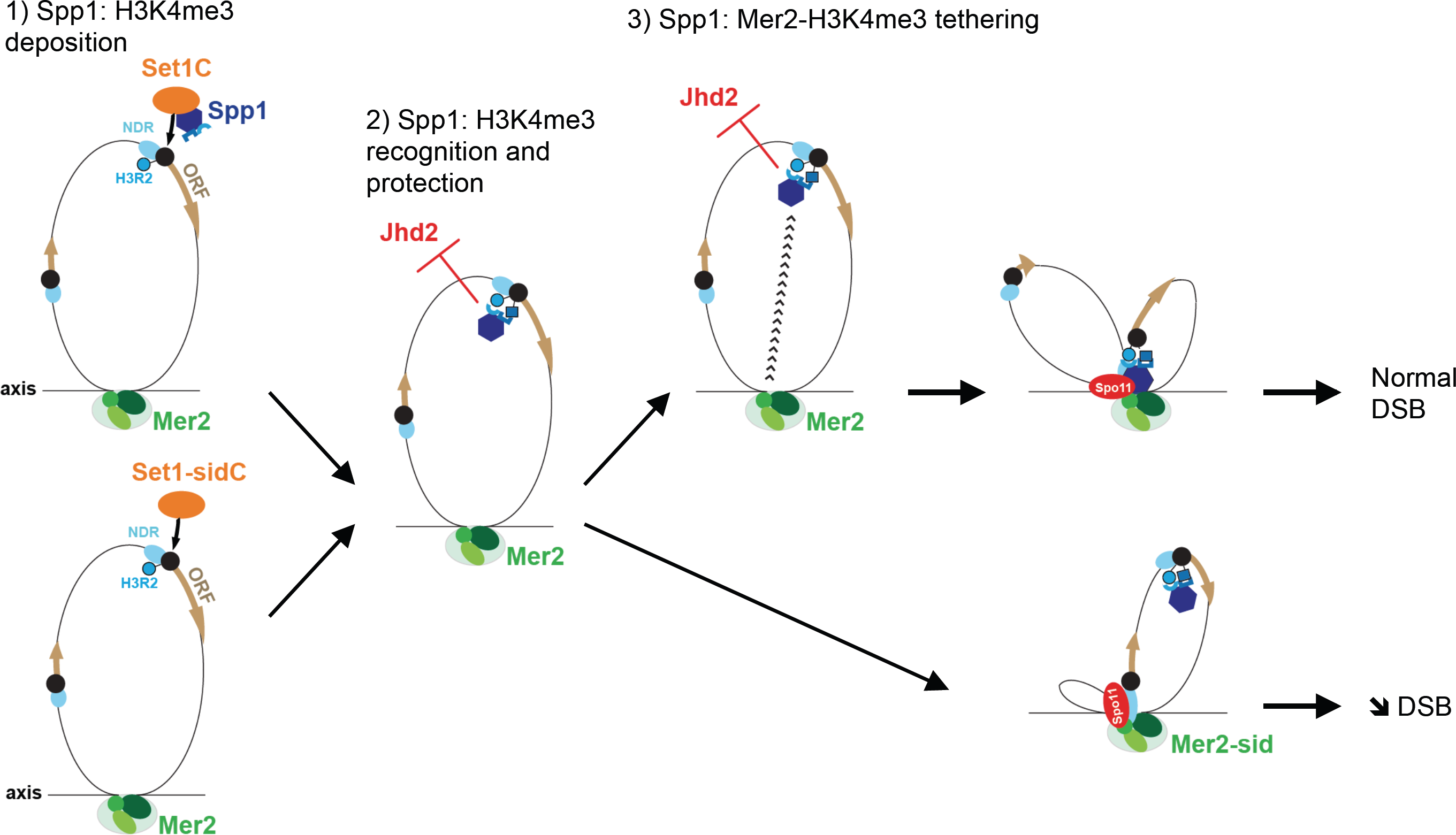
Illustration of the different functions of Spp1 for H3K4me3 and meiotic DSB formation. 1) in the Set1 complex, Spp1 has a role to allow catalysis of H3K4 trimethylation by Set1, but this function is not essential, since the *set1-sid* mutant still maintains high levels of H3K4me3. 2) In addition, Spp1 maintains H3K4me3 levels, not by stimulating Set1 catalytic activity, but likely by binding H3K4me3 with its PHD finger. This can take place without interaction with the Set1 complex. We propose this may protect H3K4me3 from active demethylation, by the Jhd2 enzyme. 3) Finally, the simultaneous binding of Spp1 to H3K4me3 and to the axis-associated Mer2 protein is essential to promote efficient DSB formation by Spo11. It has to be noted that a PHD finger mutant of Spp1 (W45A) is still able to bind Mer2 (17), so recognition of H3K4me3 by Spp1 PHD finger is not a prerequisite for its subsequent binding to Mer2. NDR: nucleosome-depleted region; Black circle: first nucleosome of genes; H3R2: arginine 2 of histone H3, in its non-asymmetrically methylated form. The blue square represents H3K4me3.

The closest homolog of Spp1 in Mammals, CXXC1 (CFP1) has a N-terminal PHD finger that is able to interact with histone H3K4 methylated peptide *in vitro*, and seems important to allow Set1 complex binding to chromatin *in vitro* (37), However, the *in vivo* function of this PHD finger has not been determined. In addition, CXXC1 contains a motif that binds unmethylated CpG islands, allowing CXXC1 to restrict H3K4 methylation by the Set1 complex to these sites at promoters (38, 39). It is not known if CXXC1 may associate with these regions without the associated Set1 complex. Finally, CXXC1 was reported to interact with the DNMT1 methyltransferase, through a region distinct from its Set1 interacting region (40). However it was not investigated if *in vivo* this interaction occurs when CXXC1 is in the Set1 complex. To conclude, so far no Set1 complex independent function in H3K4me3 levels have been described for the homolog of Spp1 in mammals.

### Why using the same protein to regulate deposition and read H3K4me3, and not another PHD finger protein?

Although Spp1 is required both for deposition, reading of H3K4me3 and tethering to DSB proteins, we show here that surprisingly, DSB formation is as proficient when Spp1 is physically separated from the Set1 complex as in the wild-type conditions. One could thus wonder why Spp1 divided its tasks in meiosis to both participate to H3K4me3 deposition and work as a reader and tether of this mark. There are 18 other PHD finger proteins in *S. cerevisiae*, among which 8 are able to bind H3K4me2/3 *in vitro* (28). Our findings indicate that any PHD-finger motif that could interact with Mer2 would be able to promote DSB formation as Spp1. Maybe it is just by chance that evolution selected the same protein for fulfilling its complementary functions. However, the fact that this configuration may also be conserved in mammals makes it unlikely (see below). Since we revealed here that Spp1 is able to bind H3K4me3 independently of the Set1 complex, it is possible that in wild-type cells, Spp1 “persists” on H3K4me3 after the passage of the Set1 complex, and this constitutes a first step in the tethering of DSB sites to the chromosome axis (Figure 7). There may thus be an evolutionary advantage of using the same protein in the three sequential complexes without these complexes having to be physically linked.

### H3K4me3 function for meiotic recombination is not related to a change in chromatin accessibility but solely to anchor meiotic DSB sites to the chromosome axis

It could be argued that H3K4me3 could only be a sign of open chromatin regions, and if it had any role, it could just be to make chromatin more accessible for DSB formation. Our present data argue against this. Indeed, in our *mer2_sid* mutant, the Set1 complex is intact, and thus chromatin modifications are unaltered compared to wild-type. Nevertheless, the DSB phenotype recapitulates the one seen in the absence of H3K4 methylation or Spp1, demonstrating that impaired tethering of DSB sites to the chromosome axis is the only cause of the DSB reduction and redistribution and that open chromatin structure is not sufficient for normal DSB formation.

### Conservation of Spp1 functions in other organisms and unifying role of H3K4me3 or CpG islands at meiotic recombination sites

Outside of budding yeast, histone H3K4me3 is emerging as being widely used for meiotic recombination. Two main pathways for choosing sites of meiotic recombination have been identified (41). The first one, which occurs in mice, humans, cattle and a certain number of other vertebrates involves PRDM9, which promotes both H3K4me3 and meiotic DSB formation at its binding sites (11-14). The other one uses open chromatin at promoters, enriched in H3K4me3 and/or CpG islands, to direct meiotic recombination (5, 8-10, 42). Although the role of H3K4me3 in meiotic recombination has not been proven apart from in budding yeast, interesting findings showed recently that CXXC1, the homolog of Spp1 in mammals, is able to interact in two-hybrid tests both with PRDM9 and IHO1, the proposed functional homolog of Mer2 (3, 43, 44). Like Spp1, CXXC1 is a member of the Set1 complex, and has a PHD finger that may recognize H3K4me3. In addition, it also binds unmethylated CpG at promoters. A model was proposed in which PRDM9, by interacting with CXXC1, would direct it away from promoters, and would tether the PRDM9-bound sites to the axis for Spo11 cleavage. In *Prdm9-/-* mice or in organisms that do not have PRDM9, CXXC1 would bind H3K4me3 and/or CpG islands and tether theses sites to chromosome axis-bound IHO1, exactly as in budding yeast. Since we have shown that in budding yeast, Spp1 is able to bind H3K4me3 outside of the Set1 complex, CXXC1 may similarly recognize H3K4me3 deposited by PRDM9 outside of the Set1 complex with which it is normally associated. This likely conservation is fascinating, and indicates that H3K4me3 and/or CpG islands could be an evolutionary conserved tether for meiotic recombination sites.

## Methods

### Yeast manipulations

All yeast strains are derivatives of the SK1 background and are listed in Supplemental Table S2. For synchronous meiosis, cells were grown in SPS presporulation medium and transferred to 1% potassium acetate with vigorous shaking at 30°C as described (45). For strain constructions and spore viability measurements, sporulation was performed on solid sporulation medium for two days at 30°C.

### Yeast strains constructions

Yeast strains were obtained by direct transformation or crossing to get the desired genotype. All transformants were confirmed using PCR discriminating between correct and incorrect integrations and sequencing for epitope tag insertion or mutagenesis.

Spp1 and Mer2 were fused at their C-terminus with a TAP-tag at their endogenous locus using the pBS1539 plasmid (46). Set1 was tagged with 6 copies of HA at its N-terminus at its endogenous locus by using plasmid pOM10 (47) and Cre-Lox excision of the marker between *SET1* promoter and the tag. Swd1 was tagged at its C-terminus at its endogenous locus by 3 copies of HA (48). Site directed mutagenesis was introduced by PCR. The mutagenic PCR was performed on the region of interest where the gene was flanked by a selectable marker and transformed into yeast. For making the *spp1W45A* and *spp1Δ263-266* mutants without tag, we first introduced an HphMX drug resistant cassette behind the 3’UTR of *SPP1*. This construct was fully functional for meiotic DSB formation (S4 Fig). Next, we used this construct to introduce the desired mutation by PCR and transformation of the fragment containing the mutated gene and its 3’UTR HphMX cassette. For *mer2_sid-Flag* we used genomic DNA from a strain containing the *MER2-FLAG-KanMX* allele for PCR mutagenesis. For introducing the *set1_sid* and the *mer2_sid* mutations without an associated tag or marker, we first deleted the *SET1* or *MER2* gene with KanMX cassette by yeast transformation. We next used CRISPR-Cas9 mediated cleavage, using a plasmid encoding Cas9 and expressing a guide RNA targeted to the KanMX cassette (plasmid generously provided by G. Zhao and B. Futcher), co-transformed together with a healing *set1* or *mer2* fragment containing the desired mutation.

### Two hybrid assays

Yeast two-hybrid assays were performed exactly as described (49). *SET1, SHG1,* SPP1 and *SWD2* ORFs were PCR-amplified from SK1 genomic DNA. *MER2* cDNA sequence was amplified from pCA5-MER2 containing *MER2* cDNA, given by Scott Keeney (29).

### Southern blot to monitor DSB formation

Cells bearing the *dmc1Δ* mutation to accumulate DSBs were harvested from meiotic time courses at each time point. Genomic DNA was prepared in low melting temperature agarose plugs and digested with the *Afl*II restriction enzyme as described (45). Southern blotting and signal quantification was performed as described (17). Probes used were from nt 123046 to 124295 chr1 for *CYS3* and *DEP1* DSBs, and from nt 194848 to 196286, chr6 for *PES4* DSB.

### Tap tag purification from meiotic cells

Spp1-TAP or Mer2-TAP purification (strain VBD1266 for Spp1-TAP, VBD1877 for Spp1-TAP *set1_sid*, VBD1402 for Mer2-TAP or ORD7339 for the untagged control) was performed from 1L (2.10^10^ cells) of a synchronous meiotic culture at t = 3.5 hr in meiosis. Each purification was performed in parallel with a control untagged strain, ORD7339. The protocol was essentially as described in (46) with the following modifications: PMSF to a final concentration of 1 mM was added to the culture prior harvesting the cells. Cells were washed with TAP lysis buffer containing 1 mM PMSF resuspended in about 2 ml of the same buffer and frozen as noodles in liquid nitrogen. For lysis, cells were ground in a mortar in liquid nitrogen, and lysis was performed in TAP Lysis buffer plus PMSF 1 mM and 1X Complete EDTA-free protease inhibitor cocktail (Roche).

### Mass spectrometry analysis

Protein preparation for mass spectrometry-based proteomic analyses was as described (50). Briefly, extracted proteins were stacked in the top of a SDS-PAGE gel (NuPAGE 4-12%, Invitrogen) before in-gel digestion using trypsin (Promega, sequencing grade). Resulting peptides were analysed by online nanoLC-MS/MS (UltiMate 3000 coupled to LTQ-Orbitrap Velos Pro and Ultimate 3000 RSLCnano coupled to Q-Exactive Plus, Thermo Scientific, for Spp1 and Mer2 interactomes analyses, respectively) using a 120-min gradient. Peptides and proteins were identified and quantified using MaxQuant (version 1.5.8.3 (51)) and SwissProt database (June 2017 version, *Saccharomyces cerevisiae* S288c taxonomy). Only proteins identified with a minimum of two unique + razor peptides were taken into account for further analyses. Statistical analyses were performed using ProStaR (52). Proteins identified in the reverse and contaminant databases, proteins only identified by site and proteins exhibiting less than 3 intensity values in one condition were discarded from the list. After log_2_ transformation, intensity values were normalized by median centering before missing value imputation (replacing missing values by the 2.5 percentile value of each column); statistical testing was conducted using *limma* t-test. Differentially recovered proteins were sorted out using a log_2_(fold change) cut-off of 7 and a FDR threshold on remaining p-values of 1% using the Benjamini-Hochberg method.

### Co-immunoprecipitation and western blot

6.10^8^ cells were washed with PBS, and lyzed in 1.5 ml lysis buffer (20 mM HEPES/KOH pH7.5; 150 mM NaCl; 0.5 % Triton X-100; 10 % Glycerol; 1 mM MgCl2; 2 mM EDTA; 1 mM PMSF; 1X Complete Mini EDTA-Free (Roche); 1X PhosSTOP (Roche)) and 125U/mL benzonase (Sigma) and glass beads three times for 30 s in a Fastprep instrument (MP Biomedicals). The lysate was incubated 1 hr at 4°C with 125U/ml benzonase, and cleared by centrifugation at 13,000 g for 5 minutes. For Spp1-TAP purifications, magnetic PanMouse IgG beads (Life Technologies) were added and incubated overnight at 4°C. Beads were washed 4 times with lysis buffer and precipitated proteins were eluted by Tev cleavage (2 μg in 20 mM Tris pH8; 150 mM NaCl; 0.1 % NP-40; 5 % glycerol; 1 mM MgCl2; 0.5 mM EDTA; 1mM DTT for 1 hr at room temperature). One volume of 2 x SDS protein sample buffer was added and sample were denatured 10 min at 95°C before electrophoresis. For Swd1-HA IP, 25 μl of Protein G magnetic beads (New England Biolabs) and 5 μg of mouse HA monoclonal antibody 16B12 (Covance)) were added. After overnight incubation at 4°C, beads were washed 4 times with lysis buffer and resuspended in 30 μl of 2 x SDS protein sample buffer. The beads were heated at 95 °C for 10 min. For Mer2-Flag IP, Sigma Anti-FLAG M2 magnetic beads were added to the lysate and incubated overnight at 4°C. Beads were washed twice with lysis buffer, and eluted for 2 hrs at 4°C with 12.5 μg of Flag peptide in elution buffer (20 mM Tris pH8; 150 mM NaCl; 0.1% Tween; 10% Glycerol; 5mM MgCl2; 0.5 mM EDTA). One volume of 2 x SDS protein sample buffer was added and sample were denatured 10 min at 95°C before electrophoresis. Protein eluates were loaded onto a 4-12% SDS-polyacrylamide gel and blotted to PVDF membrane. Antibodies used were as follows: anti-H3K4me3 (MC315, Millipore, 1/5000); anti-H3K4me2 (07-030, Millipore, 1/5000); anti-PGK1 (Invitrogen, 1:20000); anti-HA (Roche, 12CA5, 1/750); anti-Myc (Santa Cruz, 9E10, 1/500); anti-Flag (Sigma, 1/1000); anti-TAP (Invitrogen, 1/2000); anti-Spp1 (rabbit polyclonal, 1/2000). The Spp1 polyclonal antibody was made in rabbit against the full-length Spp1 fused at its N-terminus with a 6His-MBP tag. Signal was detected using the SuperSignal™ West Pico Chemiluminescent Substrate (ThermoFisher) and a Chemidoc touch system (Biorad). Signal was quantified using Image J software.

### ChIP-qPCR and ChIP-chip

For each meiotic time point, 2.10^8^ cells were processed as described (49). For HA-Set1, we used 1 μg of monoclonal 16B12 anti-HA antibody (Covance) and 50 μL PanMouse IgG magnetic beads (Life Technologies). For H3K4me3, we used 2 μl of MC315 H3K4me3 monoclonal antibody (Millipore) and 30 μL Protein G magnetic beads (New England Biolabs). For Spp1-Myc, in ChIPchip experiment we used 0.8 μg of c-Myc monoclonal antibody (9E10, Santa Cruz) and 30 μL Protein G magnetic beads, and for ChIP qPCR, we used 1.6 μg of c-Myc monoclonal antibody (9E10, Santa Cruz) and 50 μL PanMouse IgG magnetic beads. Quantitative PCR was performed from the immunoprecipitated DNA or the whole-cell extract using a 7900HT Fast Real-Time PCR System (Applied Biosystems) and SYBR Green PCR master mix (Applied Biosystems) as described (5). Primers for *ADH1*, *BUD23*, *CEN13,* axis and control site have been described (17, 53). Primers for *ACS1* and *CIT2* genes amplified fragments with the coordinates: chr I, nt 44778-44843 and chr III, nt 122043-122105, respectively. Results were expressed as % of DNA in the total input present in the immunoprecipitated sample and normalized to the negative control site in the middle of *NFT1*, a 3.5 kb long gene (indicated “control site” in the figures). For microarray hybridizations, whole-cell extract or immunoprecipitated DNA was amplified and labeled with either Cy3 (whole-cell extract) or Cy5 (immunoprecipitated sample) and hybridized on an Agilent 44K yeast whole-genome oligonucleotide array as described (5). Microarray images were read using an Axon 4000B scanner and analyzed using GenePix Pro 6.0 software (Axon Instruments, Inc.). Files were converted to text files and analyzed using the R software. The signal was normalized, smoothed and peak calling was done as described before (17). To each probe of the array, the mRNA level of the corresponding gene determined from exponential cells (54) or cells at 4 hr in meiosis (55) was attributed. For probes lying in a promoter, the mRNA lvel of the downstream gene was attributed, and for probes in divergent promoters, the mean value of the two divergent genes was attributed.

### Accession Numbers

The ChIP-chip data generated in this study have been deposited in the Gene Expression Omnibus database, accession number GSE102790. Processed data for all chromosomes are provided in S3 Table.

The mass spectrometry proteomics data have been deposited to the ProteomeXchange Consortium via the PRIDE partner repository (56) with the dataset identifier PXD00xxx.

## Acknowledgments

We thank Scott Keeney and Bruce Futcher for reagents. We thank Corinne Grey and Arnaud De Muyt for critical reading of the manuscript, and Daniel Holoch and Raphaël Margueron for discussions. We thank the recombinant proteins platform of Institut Curie for the Spp1 antibody, and the support of the discovery platform and informatics group at EDyP.

